# Kinetic Modeling of mant-ATP Turnover to Interpret the Biochemically Defined Myosin SRX State

**DOI:** 10.1101/2025.08.06.667936

**Authors:** Filip Ježek, Seungyeon Julia Han, Alison S. Vander Roest, Daniel A. Beard

## Abstract

The fluorescent ATP analog mant-ATP has become a valuable tool for quantifying occupancy of the myosin super-relaxed (SRX) state, a biochemically inactive state of myosin in striated muscle. Interpretation of mant-ATP fluorescence decay kinetics is confounded by inconsistencies in state definitions and kinetic assumptions. Here, we develop a mass-action kinetic model of myosin cross-bridge cycling and mant-ATP turnover to reconcile these discrepancies and provide a mechanistic framework for interpreting SRX measurements. Our model simulates ATP label-chase experiments and demonstrates that conventional double-exponential fitting methods do not directly quantify SRX occupancy. Instead, we show that slow and fast decay phases of mant-ATP fluorescence arise from label redistribution among kinetically distinct states, not state populations in equilibrium. The model resolves several apparent paradoxes identified in recent studies by reproducing experimental observations without requiring SRX and DRX kinetic isolation or implausible equilibrium constants. Simulations further quantify the impact of experimental factors—such as ADP accumulation, photobleaching, and initial rigor state occupancy—on fluorescence kinetics and SRX estimates. These results support a revised framework for SRX quantification and suggest that label-chase experiments must be interpreted using mechanistic models to accurately assess myosin state distributions and transition kinetics.

**SIGNIFICANCE:** The relative occupancy of myosin in the super-relaxed state (SRX) is a key determinant of basal ATP turnover in muscle. Measurement of ATP exchange using the fluorescent analog mant-ATP is used to assess the relative population of myosin in the SRX versus the disordered relaxed (DRX) state. Existing approaches to analyzing data from these experiments use double exponential fits to represent the mant-ATP decay in the chase-phase of the experiment. However, quantitative and mechanistic interpretations based on these analyses remain ambiguous. We present a mechanistic model of myosin ATP turnover that reproduces observed fluorescence decays under defined conditions. The results indicate that conventional interpretations, while qualitatively reasonable, are quantitatively inconsistent, as the observed slow and fast phases are predicted to arise from ligand redistribution rather than distinct equilibrium states. This framework enables rigorous, model-based interpretation of mant-ATP assays to help clarify how myosin kinetics are reflected in mant-ATP loading-chase experiments and experimental conditions may influence apparent SRX kinetics.

## INTRODUCTION

Experiments using the fluorescent ATP analog 2’/3’-O-(N-Methyl-anthraniloyl)-adenosine-5’-triphosphate (mant-ATP) have been used since 2010 to interrogate the biochemical state of myosin in intact muscles. Stewart et al. (1) introduced the technique of following the kinetics of mant-ATP binding and unbinding to quantify the proportion of myosin in the super-relaxed (SRX) state in a rabbit skeletal muscle preparation. Together with the experiments of Hooijman et al. (2), which were the first to apply this approach to cardiac muscle, these studies were instrumental in the identification of the SRX as a distinct biochemical state of myosin. While a consensus has emerged on the existence of the SRX state, its structural basis–—particularly its association with the interacting-heads motif (IHM) conformation—remains debated (3). Craig et al. have suggested that the biochemically defined SRX state is strongly related or even equivalent to the structural myosin conformation of the interacting head motif (IHM) (4, 5). Others suggest that the structurally defined IHM and biochemically defined SRX states do not necessarily reflect functionally related states and that these states are rarely established simultaneously (3, 6) Thus, it remains an open question as to whether or not the SRX population corresponds to the IHM population. Indeed, the interpretation of exactly how the observed mant-ATP kinetics reflect the biochemical/structural state of myosin is not entirely certain (6).

In a typical, microscope-based mant-ATP protocol, a permeabilized muscle or myofibril isolate is first incubated for several minutes or hours in ATP-containing solution (1) and then washed with rigor buffer containing no ATP (2). Next, the muscle is incubated for several minutes with low concentration of mant-ATP (e.g., 250 μM for 2 minutes (2)). The labeled mant-ATP is then chased with a higher concentration of unlabeled ATP (usually 4 mM), while observing the fluorescence intensity. Since the mant-ATP shows substantially increased fluorescence in the bound state, the observed decay of fluorescence during the chase phase is interpreted to reflect the kinetics of mant-ATP (or mant-ADP) unbinding from myosin. Moreover, the kinetics of replacement of the labeled mant-ATP with unlabeled ATP is interpreted to reveal distinct populations of myosin, representing the SRX and the disordered relaxed (DRX) state. Specifically, the SRX state is associated with a relatively slower decay of mant-ATP fluorescence in a buffer solution containing unlabeled ATP.

The observed intensity decay *I*_*obs*_ (*t*) is normalized and fit to a double exponential function, in the form of either Equation (1) (1, 2) or Equation (2) (6, 7)):

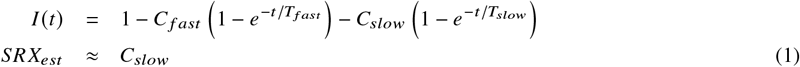

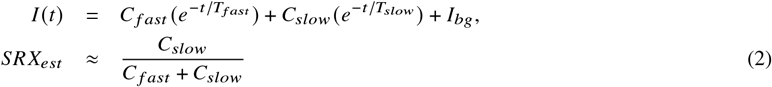

where *I*_*bg*_ is a background intensity constant. The relative occupancy of the SRX state, *SRX*_*est*_, is estimated as the amplitude of the slower phase (Equation (1) in Hooijman et al. (2)) or proportional to the slower phase (Eq. 2, as in e.g. Walklate et al. (7)), with *T*_*slow*_ ≫ *T* _*fast*_ and *T*_*slow*_ typically > 100 s. For example, Hooijman et al. report *T*_*slow*_ = 144 ± 10s in cardiac muscle (2).

Mohran et al. (6) recently highlighted three critical paradoxes associated with the interpretation of these experiments. The first is that if the SRX and DRX states are maintained in thermodynamic equilibrium, and are able to freely interconvert on a time scale faster than the replacement of mant-ATP by unlabeled ATP in the chase phase of the experiment, then the bound mant decay would follow a single exponential. Two distinct decay processes with different time scales would follow only if the SRX and DRX states are “kinetically isolated” on the timescale of the experiment, the two exponential decay function representing the proportions of myosin in the SRX and DRX states. Second, Mohran et al. point out that steady-state ATPase assays often disagree with SRX population estimates. For example, Gollapudi et al. observed that the myosin inhibitor mavacamten inhibits ATPase activity by only 70% while supposedly increasing the SRX population to effectively 100%—–a mismatch that challenges the biochemical definition of SRX (8, 9). A third paradox follows from observations of Walklate et al. (7), who report that SRX formation becomes detectable after only 200 ms of incubation with mant-ATP, yet seemingly requires hundreds of seconds to decay. This observation suggests an equilibrium mass action ratio that would overwhelmingly favor the SRX state, in conflict with interpretation of data using Equation (1) or Equation (2). The relatively fast loading of mant-ATP into the SRX state is also in apparent conflict with the requirement that the SRX and DRX states remain “kinetically isolated” on the timescale of the mant-ATP fluorescence decay.

To help resolve these paradoxes we have developed a kinetic model for mant-ATP tracer label kinetics based on a simple multi-state model of the cross-bridge cycle and SRX/DRX transitions. The ultimate goal is to develop a kinetic model that integrates the biochemical, structural, and functional observations to resolve inconsistencies and provide a unified mechanistic framework for quantification of SRX state kinetics. It is demonstrated that the three paradoxes of Mohran et al. are resolved based on a simple kinetic model and subsequently shown how the SRX prediction can be affected by experimental conditions. Specifically, model analysis demonstrates a mechanistic correspondence between the magnitude of a slower versus faster decay phase as assessed by fitting the fluorescence decay with a slow and fast phase of a double-exponential – obtained by either Equation (1) or Equation (2) – and the ATP cycling in the kinetic model. In addition, the proposed model facilitates simultaneous estimation of SRX and DRX occupancies as well as transition rate constants from mant-ATP label chase data. Furthermore, by incorporating experimental artifacts such as incomplete mant-ATP incubation or photobleaching into the fitting process, the model provides a more accurate and mechanistically grounded quantification of SRX kinetics.

## METHODS

### Computational Model

Our analysis is based on the simple kinetic model illustrated in Figure 1, similar to models of the myosin-actin cross-bridge cycle developed by this and other groups (10–13). The model assumes that myosin exists in SRX states (labeled *S*_*D*_ and *S*_*T*_), DRX states (labeled *D*_*T*_ and *D*_*D*_), or a ratcheted, rigor-like state (*R*) with no bound nucleotide. The ATP-bound DRX state *D*_*T*_ can transition into ATP-bound SRX (*S*_*T*_) with a rate proportional to first-order rate constant 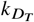, Bound ATP is hydrolyzed with rate constant *k* _−*H*_, which determines the rates of transition from *D*_*T*_ to *D*_*D*_ and from *S*_*T*_ to *S*_*D*_. The model also assumes a reverse recombination rate of *k* _*H*_, as assessed by (14). Under the unloaded calcium-free experimental conditions analyzed here ADP/ATP exchange is assumed to occur via the transition *D*_*D*_ → *R* → *D*_*T*_. The first step in this exchange, governed by rate contant *k*_leak_, involves spontaneous release of the bound ADP ·Pi and results in a ratcheted state *R*, from which the myosin needs to bind a fresh ATP (or mant-ATP) molecule. As a simplifying assumption the state *R* lumps together a strongly attached myosin state (possibly in rigor prior to mant-ATP incubation) and weakly or not attached state (possibly relaxed in ATP/mant-ATP buffer) and simply assumes that the *R* state does not have any nucleotide bound. While the transition from *D*_*T*_ to *D*_*D*_ is part of the canonical cross-bridge cycle and is typically found to be relatively rapid (associated with a time constant of less than a second), combining a futile ratcheting with ATP reattachment represents a net ATP turnover that is not associated with any cross-bridge attachment, force development, or work. This parasitic, energy-depleting futile cycle is expected to be relatively slow—operating on a time scale of many seconds. In previous interpretations of the mant-ATP assay this pathway was known as the DRX ATP turnover (6). By kinetically linking the SRX with DRX, our model does not consider spontaneous loss of ADP and/or Pi from the SRX states. Instead, we hypothesize this futile cycle is enough to explain both the incubation and chase phases of the experiments simulated below, where the labeled mant-ATP fluorophore is eventually replaced by the unlabeled ATP in the chase phase.

**Figure 1.**
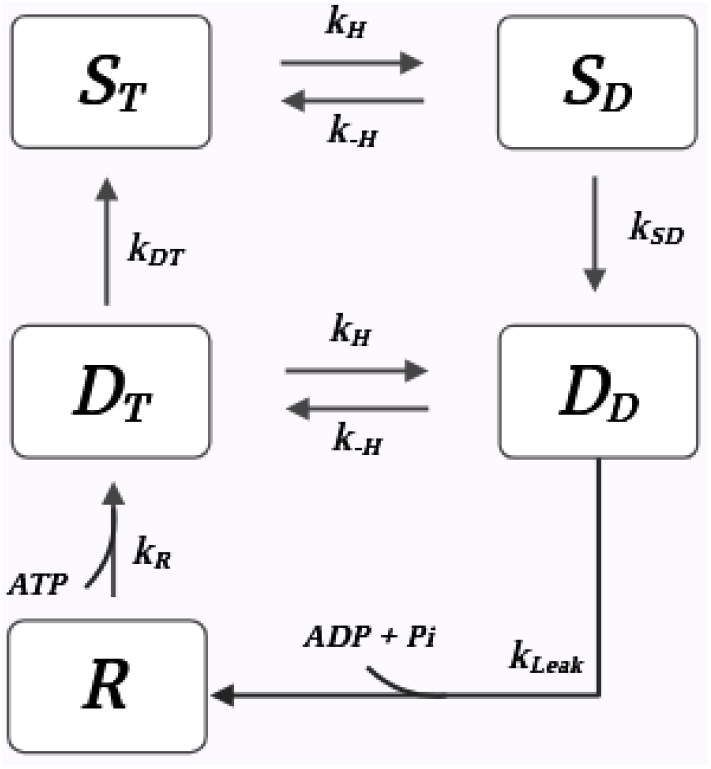
Model schematic. *S*_*T*_ – ATP-bound myosin in super-relaxed state, *S*_*D*_ – hydrolyzed ADP ·Pi-bound myosin in super-relaxed state, *D*_*T*_ – ATP-bound myosin in non-super-relaxed state, *D*_*D*_ – ADP· Pi-bound myosin in non-super-relaxed state, and *R* – nucleotide dissociated ratcheted state (weakly or strongly attached or unattached)

The probability of each state (equivalent to the relative occupancy) is denoted *P*_*x*_, where *x* represents a given state:

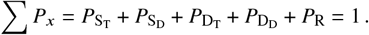

To simulate an experimental protocol where the system is initially incubated in an ATP-free rigor solution, the state probabilities are governed by the following system of equations

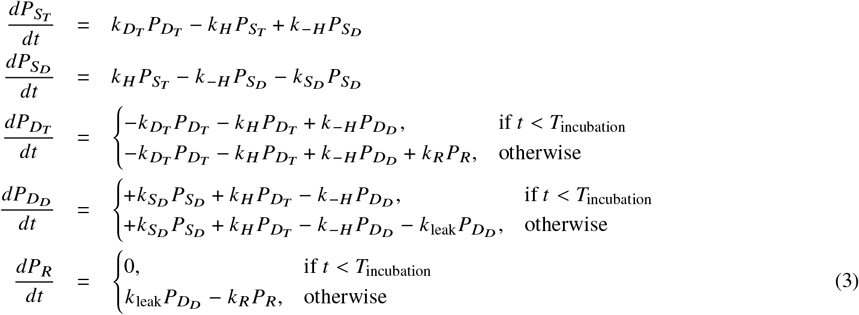

Here it is assumed that during an initial rigor phase (*t* < *T*_incubation_) there is no transition from the ratcheted state *R*, because no ATP is present. The ADP · Pi leak transition *k*_leak_ is also blocked to prevent saturation of the *R* state prior to incubation, as the population of this state is set fixed to be further analyzed.

Model analysis of the mant-ATP tracer kinetic experiment tracks the fraction of each mant-ATP/ADP-bound label in all states. We define *F*_*x*_ as the fraction of each state that has the mant label on the attached ATP/ADP, and the total probability that a myosin head has a mant label attached as *M*_*x*_ = *P*_*x*_ *F*_*x*_. From these definitions we have

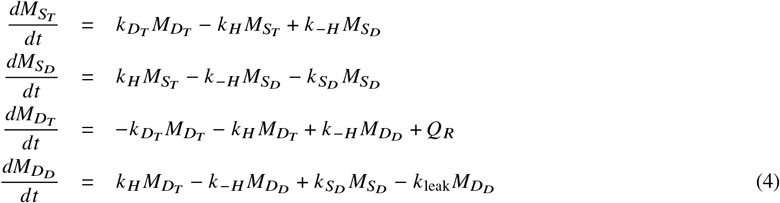

The governing equations for the mant-label, Equation (4), are coupled to the equations governing the bulk probabilities, Equation (3), via the terms *Q*_*R*_, which determine the rate at which the label is added to myosin. The rate *k* _*R*_ (*t*), associated with the transition *R* → *D*_*T*_, is the rate of addition or removal of the mant-label:

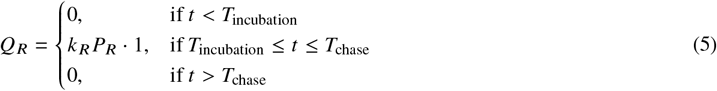

As above, we assume that during the initial rigor phase (*t* < *T*_incubation_) there is no mant-ATP present and thus *Q*_*R*_ = 0. During the incubation phase (*T*_incubation_ ≤ *t* ≤ *T*_chase_) mant-ATP is loaded onto myosin at the rate *k* _*R*_*P*_*R*_. Conversely, during the chase phase the label is lost from the system through the 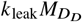.

Finally, for a direct comparison to data, simulated fluorescence intensity of the label *I*_*sim*_ is calculated as a normalized sum of *M*_*x*_ plus experimentally observed background *I*_*bg*_:

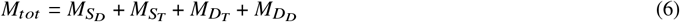

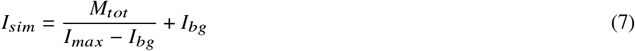

and the predicted SRX state fraction *SRX*_*pred*_ is calculated as

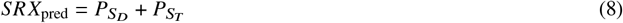

### Photobleaching

The fluorescence intensity of mant-ATP decreases over time due to degradation of the mant group by the applied excitation energy. To simulate photobleaching under conditions of steady incident light intensity the mant signal intensity is assumed to decay exponentially

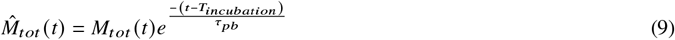

where the corrected total active mant-label 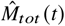 replaces the *M*_*tot*_ (*t*) in Equation (7) and *τ*_*pb*_ is a photobleaching time constant.

### ADP-Mediated Inhibition

The *k*_leak_ transition involves release of bound ADP and Pi and thus may be influenced by ADP concentration, particularly at relatively high concentrations of myosin/myofibrils, when sampled over a long incubation times and when the buffer solution does not contain any ATP regenerating system (as in (2, 7)). The potential inhibitory effect of ADP concentration (both mant-ADP and ADP) on the *D*_*D*_ → *D*_*T*_ transition is investigated by replacing *k*_leak_ in equations 3 and 4 with an effective rate constant 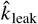, influenced by *C*_*ADP*_, the concentration of ADP in the experimental buffer:

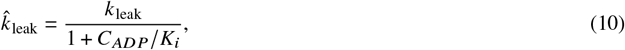

where *K*_*i*_ is an inhibition constant. The total ADP (labeled plus unlabeled) concentration is determined by the rate of turnover of the state *D*_*D*_ to *D*_*T*_:

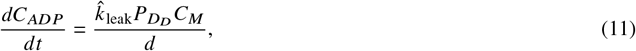

where *C*_*M*_ is the initial concentration of myosin ATPase in the experimental system and *d* is the dilution factor caused by mixing with incubation or chase solutions. In the simulated single mixing experiments, the solution with muscle is diluted by the added chase buffer of the same volume. In double mixing scenarios, the muscle specimen is diluted by the incubation buffer with mant-ATP first and then by the chase buffer, containing dark ATP. Thus,

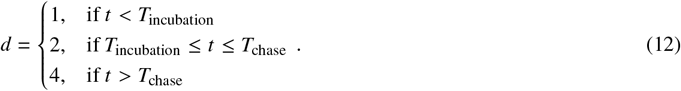

When the ADP inhibition is not considered, 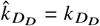.

### Experimental and Simulation Protocol

The kinetic model is applied to simulate mant-ATP label-chase experiments, such as those of Hooijiman et al. (2) and Walklate et al. (7), in which either permeabilized fibers or myofibrils are first incubated in an ATP-free rigor solution. A mant-ATP containing solution is then introduced for an incubation period, during which the mant-ATP label binds to myosin heads. The incubation period is followed by a chase period, in which unlabeled ATP is introduced and the mant label is displaced by the unlabeled ATP.

We use the convention that *t* = 0 represents the beginning of the chase phase. I.e., *T*_*chase*_ = 0 in the above equations. The mant-ATP incubation phase (also referred to as the “loading phase”, “labeling phase” or “aging time” elsewhere) is defined to begin at *t* = *T*_*incubation*_. An initial population of cross-bridges is assumed to exist in the ratcheted state *R* during the initial rigor state, *R*_0_ ∈ (0, 1). Since no ATP is present in the rigor state, the probability *P*_*R*_ remains constant at this fixed value during the rigor state. The probabilities 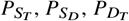, and 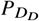 are assumed to be in equilibrium during the rigor state, with the equilibrium distribution of states is determined by the steady-state solution to Equation (3) for *t* < *T*_*incubation*_.

### Model Baseline Identification

Model parameters (see Table 2) *k* _*H*_ and *k* _*R*_ are set to *k* _*H*_ = 18 s^−1^ and *k* _*R*_ = 16s^−1^ (7, 15). Although these values for bovine masseter are used also for the Hooijman case, the match to experimental data is minimally sensitive to the values chosen. Other parameter values are estimated based on the fluorescence data of ATP chase after mant-ATP incubation experiments reported in the source literature, as specified in the table 2. For simulations of the experiments on rabbit ventricular muscle reported in Hooijman et al. (2), a pseudo-data set was constructed from Table 1 of Hooijman et al. (2), using Equation (1) to generate the chase-phase fluorescence time course. The sum of squared difference of simulated and observed fluorescent decay data ((*I*_*sim*_ − *I*_*obs*_)^2^) was minimized by optimizing *k*_leak_, 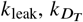, and 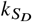. Estimated parameter values are listed in Table 2.

**Table 1.**
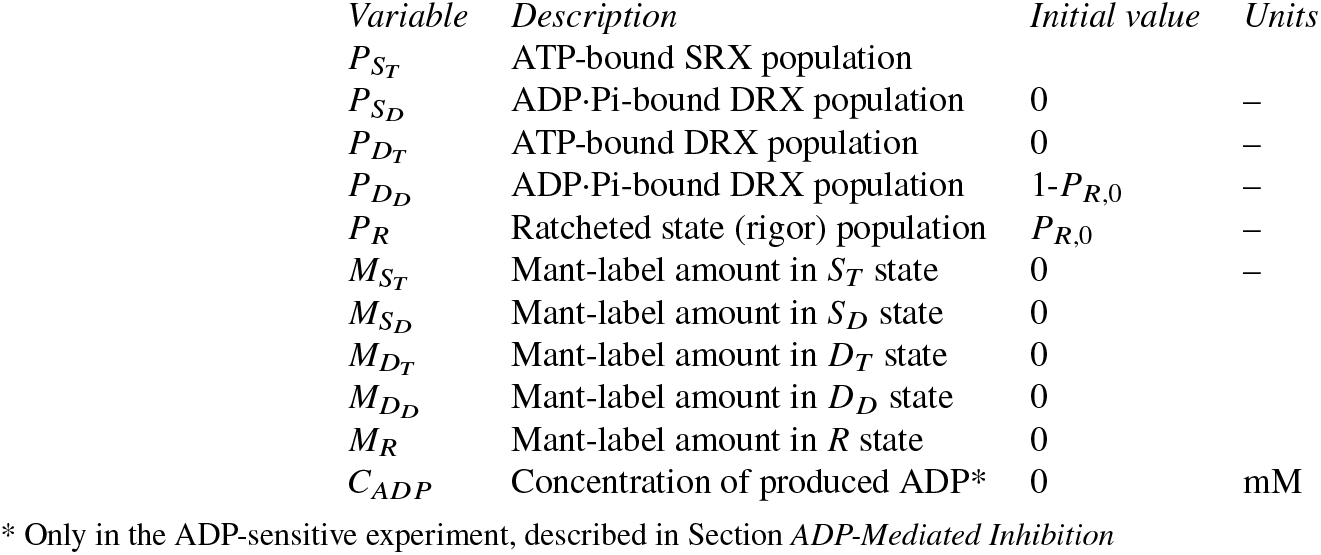
Model state variables.

**Table 2.** Model parameters for different model configurations. Values from a) Hooijman et al. (2), Table 1 for cardiac tissue (rabbit ventricle), b)Walklate et al. (7), Fig. 1, for skeletal muscle (bovine masseter), c) (15), table 1, (bovine masseter), d) as 1/10 of the forward rate (14) e) (16). Model variants RF - Rigor fraction analysis (Fig. 6), PB - photobleaching analysis (Fig. 7), AS - ADP accumulation sensitivity analysis (Fig. 5).

| Optimized parameters |  | Value |  |  |  |  | Unit |
| --- | --- | --- | --- | --- | --- | --- | --- |
| Parameter | Description | Hooijman<sup>a)</sup> | Walklate<sup>b)</sup> | Walklate<sup>b)</sup>,<br>RF | Walklate<sup>b)</sup>,<br>PB | Walklate<sup>b)</sup>,<br>AS |  |
| $k_{\text{leak}}$ | ADP·Pi release rate from $D_D$ | $74 \times 10^{-3}$ | $27 \times 10^{-3}$ | $26 \times 10^{-3}$ | $24 \times 10^{-3}$ | $48 \times 10^{-3}$ | $\text{s}^{-1}$ |
| $k_{D_T}$ | $D_T$ to SRX transition rate | $36 \times 10^{-3}$ | $25 \times 10^{-3}$ | $26 \times 10^{-3}$ | $37 \times 10^{-3}$ | $17 \times 10^{-3}$ | $\text{s}^{-1}$ |
| $k_S$ | SRX to $D_T$ transition rate | $8.1 \times 10^{-3}$ | $4.7 \times 10^{-3}$ | $4.7 \times 10^{-3}$ | $10^{-5}$ | $4.9 \times 10^{-3}$ | $\text{s}^{-1}$ |
| Fixed parameters |  |  |  |  |  |  |  |
| $k_{H^{\ddagger}}$ | ATP hydrolysis ($D_T$ to $D_D$ and $S_T$ to $S_D$) rate | $18^{\text{c)}$ | $18^{\text{c)}$ | $18^{\text{c)}$ | $18^{\text{c)}$ | $18^{\text{c)}$ | $\text{s}^{-1}$ |
| $k_{-H^{\ddagger}}$ | Reverse hydrolysis recombination ($D_D$ to $D_T$ and ($S_T$ to $S_D$)) rate | $1.8^{\text{d)}$ | $1.8^{\text{d)}$ | $1.8^{\text{d)}$ | $1.8^{\text{d)}$ | $1.8^{\text{d)}$ | $\text{s}^{-1}$ |
| $k_{R^{\ddagger}}$ | $R$ to $D_T$ transition rate, including ATP binding | $16^{\text{b)}$ | $16^{\text{b)}$ | $16^{\text{b)}$ | $16^{\text{b)}$ | $16^{\text{b)}$ | $\text{s}^{-1}$ |
| $D_i$ | Incubation duration | $120^{\text{a)}$ | $60^{\text{b)}$ | $60^{\text{b)}$ | $60^{\text{b)}$ | $60^{\text{b)}$ | s |
| $K_i$ | Inhibition constant on $k_{\text{leak}}$ transition | | | | | 0.1 | $\mu\text{M}$ |
| $T_{pb}$ | Photobleaching time constant | | | | 500 | | s |
| $P_{R,0}$ | Initial rigor fraction | $80^{\text{e)}$ | $80^{\text{e)}$ | 40 | $80^{\text{e)}$ | $80^{\text{e)}$ | % |
| $I_{max}$ | Maximal measured fluorescence intensity | $100^{\text{a)}$ | $106^{\text{b)}$ | $106^{\text{b)}$ | $106^{\text{b)}$ | $106^{\text{b)}$ | % |
| $I_{bg}$ | Maximal measured fluorescence intensity | $7^{\text{a)}$ | $99^{\text{b)}$ | $99^{\text{b)}$ | $99^{\text{b)}$ | $99^{\text{b)}$ | % |

‡ ATP-requiring (labeling) transitions
‡ Little sensitive within the tested range.

### Model Implementation

The model is implemented in the Modelica language using the Bodylight v1.1 library (https://github.com/creative-connections/bodylight.mo) and is available on GitHub at https://github.com/beards-lab/mant-ATP-SRX. Open-source tools, such as OpenModelica, can be employed for simulation. Please see the code documentation for more information.

## RESULTS

### Analysis of the Hooijman et al. Experiment

Fig. 2 shows a simulation of a microscope-based mant-ATP assay of rabbit ventricular muscle strips, reported by Hooijman et al. (2). For this experiment, the mant-ATP incubation time is 2 minutes, and the initial rigor state probability is assumed to be *R*_0_ = 80% (16). Parameter values for these simulations are listed in Table 2. Simulations in Fig. 2 do not account for photobleaching or ADP-mediated inhibition. Panel D shows optimized fit of the model to the fluorescence decay of the Hooijman et al. experiment (Fig. 2B) from Hooijman et al. (2)), demonstrating perfect match of the kinetic model to the observed data.

**Figure 2.**
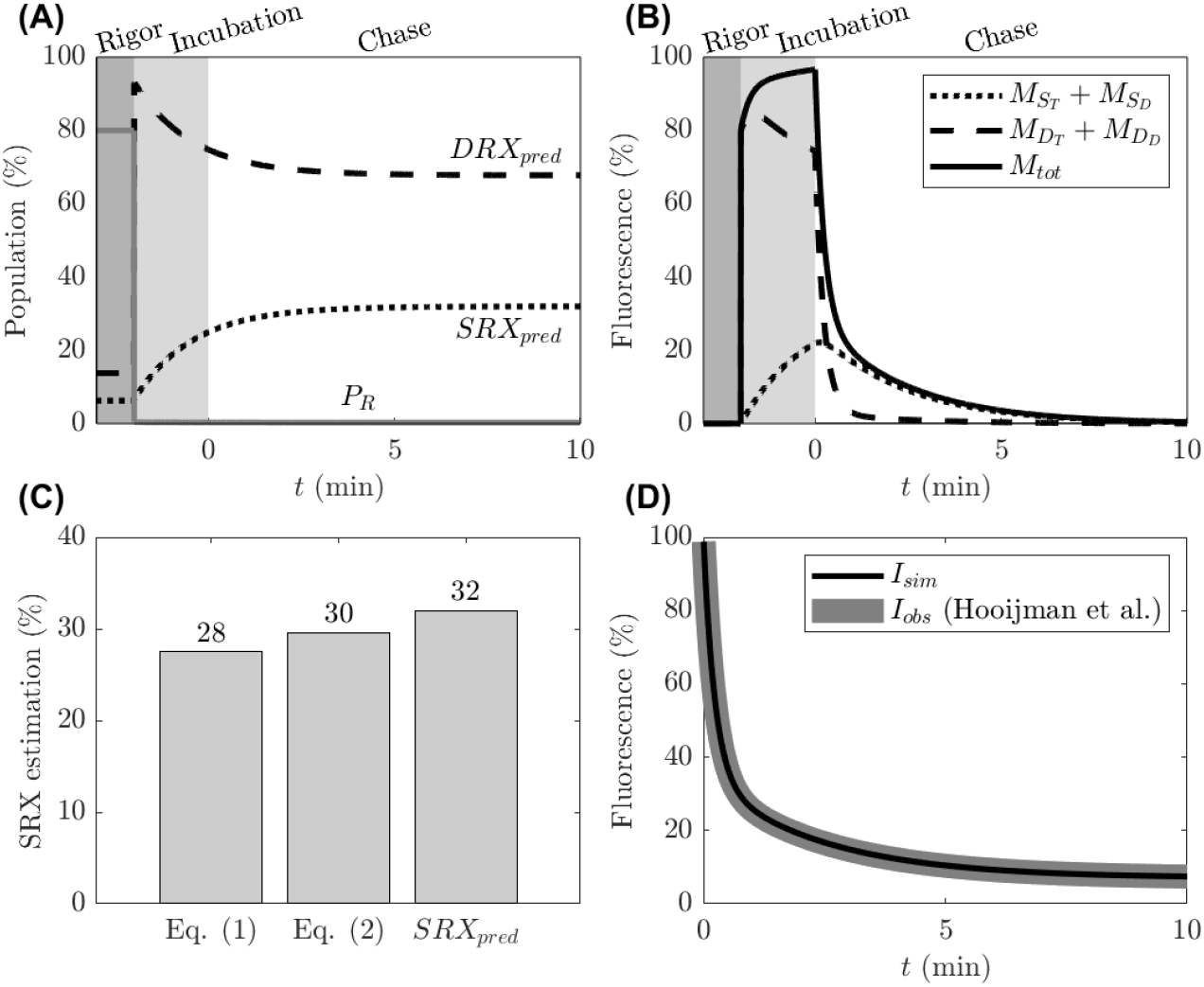
Simulation of the Hooijman et al. (2) label-chase experiment with label incubation time of 2 minutes. (A) The simulated state probabilities show redistribution from the ratcheted (*R*) and ADP-bound DRX states to the SRX states following addition of mant-ATP to initiate the label incubation phase. (B) The predicted kinetics of fluorescent labeled states shows that while the labeling reaches maximal capacity during the incubation phase, the label distribution between the DRX and SRX states has not reached steady state at the end of the incubation phase. (C) The estimated SRX fraction, based on fits of simulated data to Equation (1) is 28% and to Eq.2 (2) is 29 %, while the simulated predicted steady-state probability of both SRX states is 32%. (D) Simulated fluorescent intensity *I*_*tot*_ during the chase phase perfectly reproduces the fluorescence decay in the ATP chase phase *I*_*obs*_ observed by (2).

Panel A shows the simulated bulk state probabilities during the rigor, incubation, and chase phases of the experiment. During the incubation phase *P*_*R*_ rapidly drops from 80% to zero following the introduction of mant-ATP. The rapid drop in *P*_*R*_ is associated with a rapid spike in 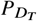 at the beginning of the incubation phase. However, the *D*_*T*_ (DRX) state mostly rapidly hydrolyzes to the *D*_*D*_ state. Thus the *DRX* states rise to about 95% at the beginning of the incubation phase. The distribution of probability into the SRX states is gradual over the two-minute incubation and ten-minute chase phase. The DRX state probability 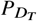 remains in dynamic equilibrium with the 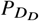 in a ratio of around 1:10 (14), yet facilitates fine balance between SRX and DRX.

Panel B shows the corresponding predicted distribution of fluorescent label states during this experiment. During the incubation phase the fluorescent label accumulates primarily in the *M*_*S*_ and *M*_*D*_ states since the proportion of the system in the 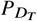 state is relatively minor. During the chase phase the decay of the 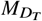 and 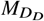 state represents the fast phase of fluorescent decay and the decay of the 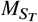 and 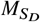 represent the slow phase. Using Equations 1 and 2 to assess the simulated data, or equivalently to assess the pseudo data (constructed from row 1, Table 1 of (2)) from Panel D, yields estimates of the slow phase fraction of 28% and 30%, as illustrated in Fig. 2C. The long-time steady-state predicted SRX value of 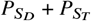 is almost the same, 32%.

### Analysis of the Walklate et al. Experiment

Model simulations fitting a sample stopped-flow experiment of Walklate et al. ((7), Fig 1D), using bovine masseter myofibrils, are illustrated in Fig. 3. The data are presented as in Walklate et al., Figure 1A, assuming normalized to maximum intensity of the loading phase. As for Fig. 2, simulations in Fig. 3 do not account for photobleaching or ADP-mediated inhibition. Panel D demonstrates an essentially exact fit of the decay label, which is associated with an estimated SRX fraction of *P*_*s*_ = 34%.

**Figure 3.**
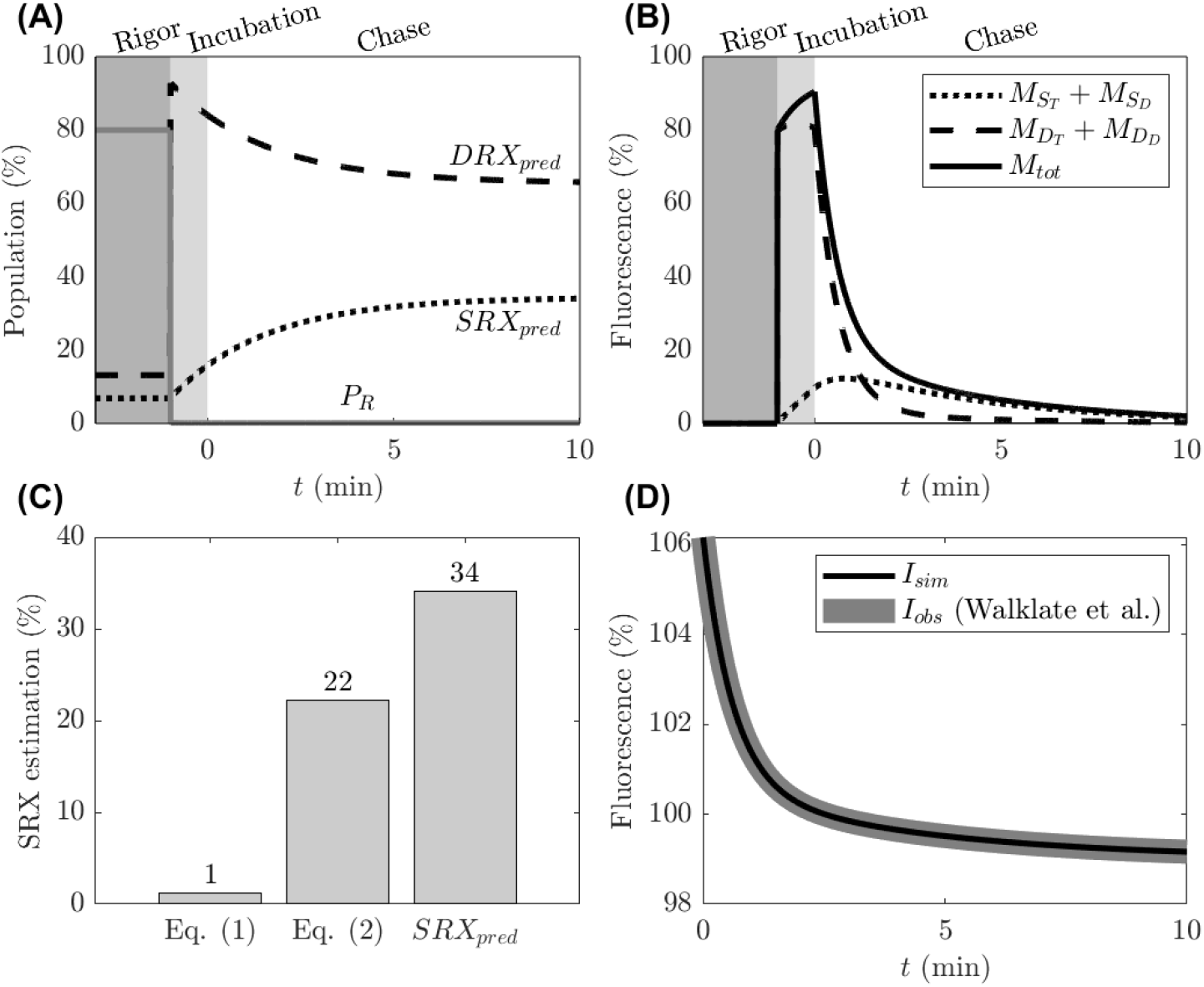
Simulation of the Walklate et al. sample experiment ((7), Fig. 1A) (A) The simulated state probabilities show transition from the ratcheted (*R*) and ADP-bound (*P*_*D*_) states to the SRX (*P*_*S*_) and ATP-bound DRX 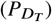 states following addition of mant-ATP to initiate the label incubation phase. Notice shorter incubation (60 seconds) compared to Fig. 2 (B) The predicted kinetics of fluorescent labeled states shows unsaturated labeling during incubation time. (C) The estimated SRX fraction obtained from fitting the simulated data is approximately 23%, whereas Equation (1) was found to be unusable without background subtraction. The simulated steady-state probability of the SRX state is 34%. (D) Simulated fluorescent intensity *I*_*tot*_ during the chase phase is fit to reproduce the fluorescence decay in the ATP chase phase *I*_*obs*_ observed by (7) Fig. 1A.

However, as reported in panel C, fits of the decay to double exponential Equation (2), yield estimate of 22%, approximately two-thirds of the value associated with the kinetic model that exactly matches the kinetic data. This discrepancy may contribute to differences in predictions of the SRX fraction made using different experimental protocols.

Walklate et al. (7) observed that the slow decay phase during the chase phase of the experiment (attributed to the SRX state) is established with relatively short (≈200 ms) incubation times. To explore this behavior we simulated a range of incubation times, from 0.2 seconds to 15 minutes, using the model parameters estimated based on Fig. 3. For each simulated incubation time, we simulated the chase phase and fluorescent decay, as in Fig. 3, and computed the estimated SRX fraction from Equation (2). Results are summarized in Fig. 4.

**Figure 4.**
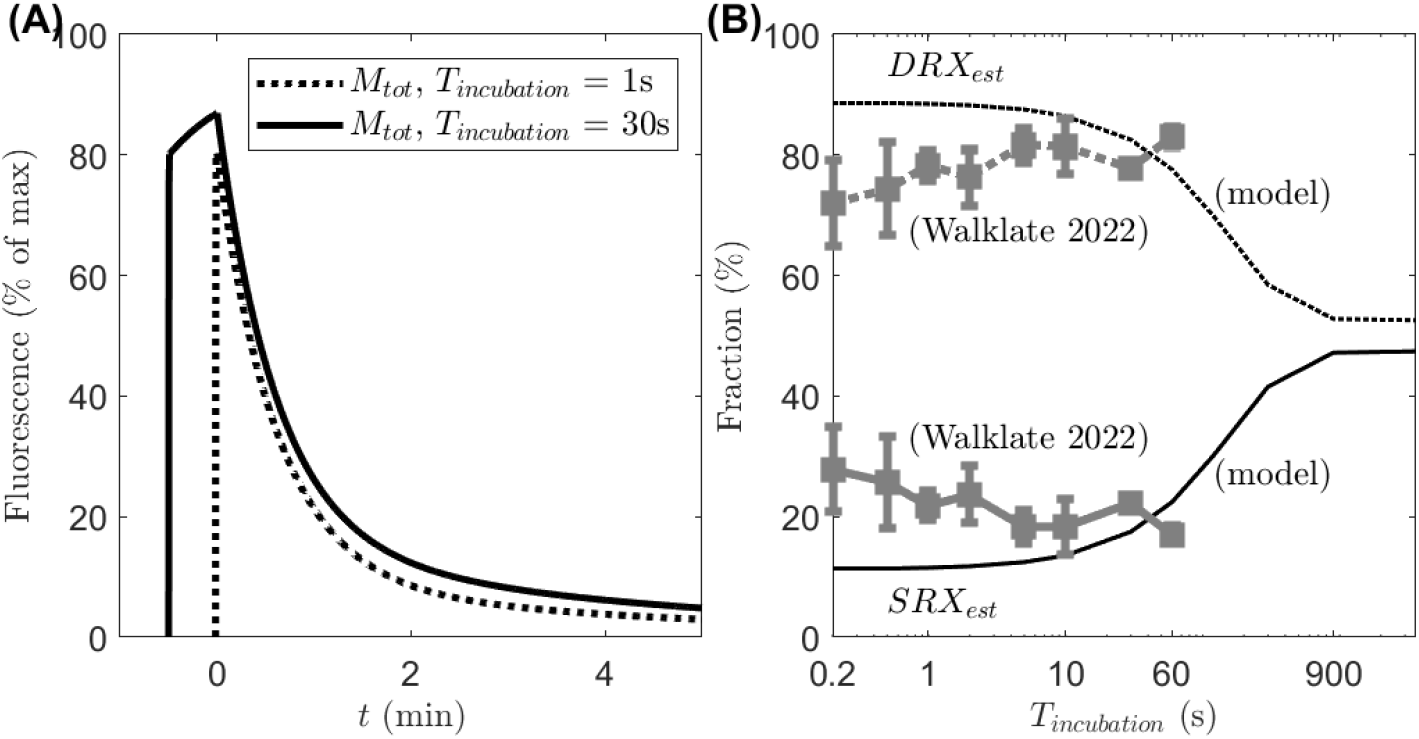
Effect of mant-ATP incubation time on estimated SRX and DRX fraction. A. Simulation of total mant label in the system for 1- and 30-second incubation time. B. Estimated SRX fraction from Equation (2) is plotted as a function of incubation time, while the predicted steady state *SRX*_pred_ is 34%, regardless of the incubation time. (Estimated DRX fraction is 100% minus the estimated SRX fraction.) Data from Walklate et al. (7) Fig. 1D are shown for comparison.

The left panel of Fig. 4 shows the predicted intensity time course for 1- and 30-second incubation times. For both incubation times, there is an initial effectively instantaneous loading of 80% of the total myosin states as the initial rigor fraction (here fixed at 80%) rapidly transitions to the *D*_*T*_ and subsequently the *D*_*D*_ state. Loading of the label slows following this initial fast phase and the magnitude of intensity reached for the 30-second incubation is only slightly greater than that for the 1-second incubation and the shapes of the decay curve during the chase phase show similar bi-exponential behavior. Estimating the SRX fraction based on fitting Equation (2) to simulated fluorescence intensity *I*_*sim*_ for the 1-second incubation simulation yields 12%, while estimating the SRX fraction based on Equation (2) for the 30-second incubation simulation yields 18%. Thus using Equation (2) to assess the SRX fraction from these simulated data sets predicts that the estimated slow phase fraction is largely independent of incubation time over this range of incubation times.

The right panel of Fig. 4 shows the results of analyzing simulated data obtained for a range of incubation times using Equation (2) to assess the SRX fraction. Simulated results are compared to the data of Walklate et al.(7). Similar to the results reported by Walklate et al.(7) the simulated analysis shows that the estimated SRX fraction remains effectively constant over a certain range of incubation times: Over the loading times of 0.2 to 10 seconds the estimated SRX fraction increases from 11% to 14%. However, the estimated SRX fraction increases to 22% for the 60 second incubation time, while the analysis of Walklate et al. (7) shows a decrease from approximately 28% at 0.2 seconds to 17% at the 60 second incubation time. At higher incubation times the estimated SRX fraction saturates at 50%. Most importantly, the true steady state SRX population *SRX*_*pred*_ = 34% for the simulated model, regardless of incubation time.

These results illustrate how the estimated slow phase fraction may be largely independent of incubation time over a wide range of incubation time, yet fully consistent with our simple mass-action kinetic model. Walklate et al report that measuring longer incubation times was limited due to mnat-ADP build-up (7). Discrepancies between simulations and data of Walklate et al. in Fig. 4B may derive in part of the effects of ADP-mediated inhibition of the *D*_*D*_→ *D*_*T*_ state transition, which are not accounted for in Fig. 4.

### Accounting for ADP-Mediated Inhibition

Walklate et al. (7) suggest that a build-up mant-ADP becomes limiting for incubation times longer than 60 seconds in their experimental setup. ADP-mediated inhibition of the *D*_*D*_ → *D*_*T*_ transition is modeled using Equation (10). Using this formulation, sample data from Walklate may be fit and analyzed using a range of values of *K*_*i*_. Fig. 5D shows an example fit to the single-mixing chase phase decay, assuming *K*_*i*_ = 0.1 μM for the 1-minute incubation experiment. Further decrease in *K*_*i*_ (that is an inhibition more sensitive to ADP) makes the model unable to fit the data anymore. Panels A and B show the simulated time courses of state probabilities and label occupancy for this fit. At this level of ADP-mediated inhibition the predicted *SRX* at the end of the chase phase is 26%—significantly lower than the 34% value observed in Fig. 3 with no ADP inhibition.

**Figure 5.**
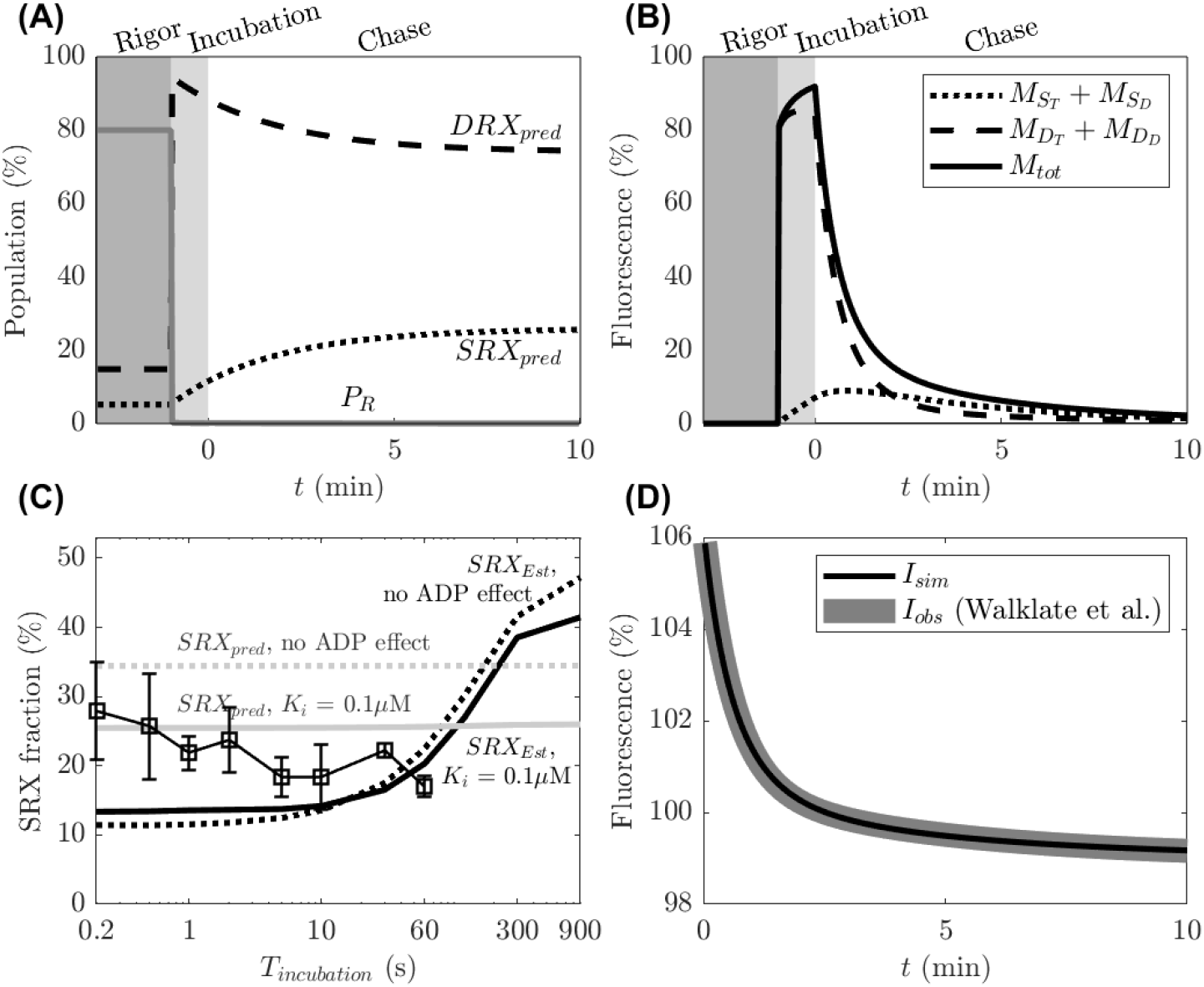
Simulation with ADP-dependent inhibition of *D*_*D*_ → *D*_*T*_ transition, with *K*_*i*_ = 0.1 μM. (A) The simulated state probabilities show that ADP-mediated inhibition leads to a lower predicted steady-state SRX occupancy (*SRX*_*pred*_ = 26% compared to 34% with no inhibition). (B) The predicted kinetics of fluorescent labeled states are equivalent to the simulated kinetics for the ADP-independent case of Figure 3. (C) The estimated SRX fraction, based on fits of simulated data to Equation (2) as a function in incubation time for the ADP inhbition case (black, dotted) is similar to the ADP-independent case (black, solid). (D) Simulated fluorescent intensity *I*_*tot*_ during the chase phase almost perfectly reproduces the fluorescence decay in the ATP chase phase *I*_*obs*_ observed by (7).

Although ADP-mediated inhibition results in a reduction of the SRX states probability, it is predicted to have little effect on the estimated slow phase fraction from the chase phase data. Fig. 5C shows the predicted effect of ADP-inhibition on SRX estimation at *K*_*i*_ = 0.1 μM as a function of incubation time for the Walklate et al. double-mixing incubation-sweep experiment. Incorporation of ADP inhibition leads to a slightly better match to the data compared to Fig. 4B. Thus, the decrease in estimated slow phase fraction observed by Walklate et al. might be explained by ADP-mediated inhibition as represented in this kinetic model. As demonstrated in Fig. 2 and Fig. 3, the slow phase fraction estimated from Equation (2) does not necessarily correspond to relative occupancy of the SRX state. Moreover, this prediction demonstrates that a kinetic mechanism that decreases the relative occupancy of the SRX state can have little effect on the slow phase fraction estimated from the double-exponential analysis of the chase phase data.

### Effect of Initial Rigor Fraction

During the muscle preparation, the muscle fibers are demembranated for several hours and stored in various solutions (protocols differ). While some studies explicitly report a pre-incubation step in rigor (ATP depleted) buffer, these details are not always reported. Simulations above assume a finite initial occupancy of the *R* rigor state before the incubation phase. Once ATP is introduced into the system, the myosin heads rapidly accumulate in the *D*_*D*_ state. In the simulations above, initial fraction of myosin population in the ratcheted state (*R*_0_), representing the initial rigor, was set to 80% (16). To evaluate the effect of this parameter on the predicted steady-state SRX, we reoptimized the model fit to the data from Fig. 1A of Walklate et al. (7) for each of *R*_0_ = (0, 20, 40, 60, 80, 100). Fig. 6D shows a fit to input data for *R*_0_ = 40%, while panels A & B show the predicted state probabilities and label occupancy for this fit. Panel C shows the predicted steady-state SRX fraction, 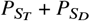, for the range of assumed *R*_0_ values. While the value of *R*_0_ has a large effect on the amount of label *M*_*tot*_ in the initial phase of the incubation (compare Fig. 3B and 6B), it has a minor observable effect on the predicted steady-state SRX fraction (Fig. 6C). In other words, all assumed values for *R*_0_ result in similar predicted SRX and model predictions in terms of SRX occupancy are relatively insensitive to *R*_0_.

**Figure 6.**
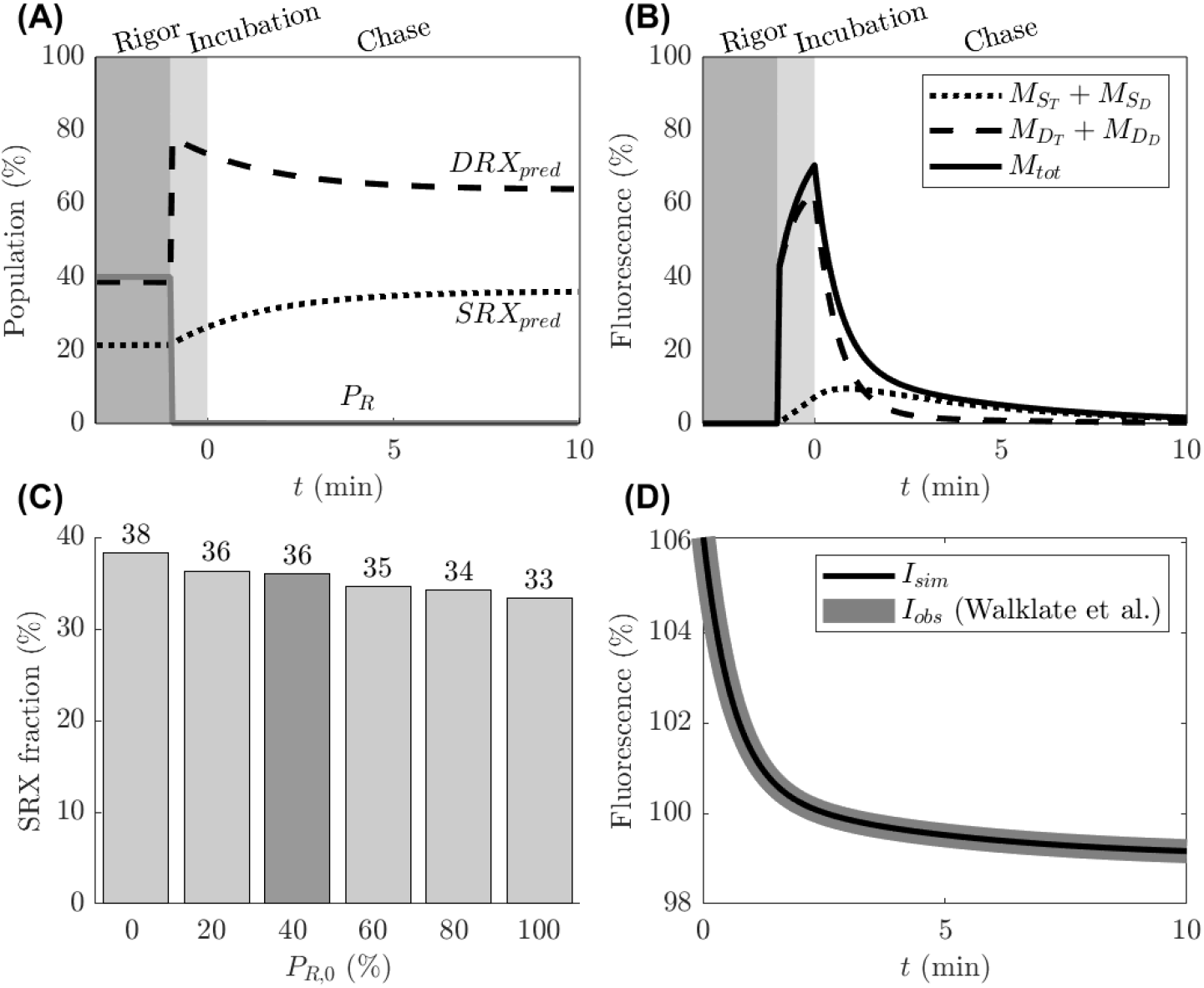
Effect of initial rigor fraction *R*_0_, decreased to 40%, on predicted steady-state SRX fraction. (A) The simulated state probabilities show transition from the ratcheted (*R*) and ADP-bound (*P*_*D*_) states to the SRX (*SRX*_*pred*_) and DRX 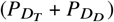 states following addition of mant-ATP to initiate the label incubation phase (B) The predicted kinetics of fluorescent labeled states. (C) Different values of initial rigor state fraction *R*_0_ yield little effect on predicted steady-state SRX fractions at lower *R*_0_. (D) Simulated fluorescent intensity *I*_*tot*_ during the chase phase is fit to reproduce the fluorescence decay in the ATP chase phase *I*_*obs*_ observed by (7) Fig. 1A using *P*_*R*,0_ = 40%.

### Accounting for Photobleaching

Photobleaching via repeated excitation can gradually reduce the mant fluorescence signal by rendering mant fluorophores unable to fluoresce. The photobleaching effect may be reduced through the experimental design (e.g. (6)) or accounted for in the data analysis (e.g. using a linear approximation, as in (17)). We evaluated the potential impact of photobleaching on predicted steady state SRX using the photobleaching model of Equation (9), which assumes a constant steady excitation and bleaching beginning at the start of the incubation phase.

Fig. 7D shows a fit of the model to the data of Walklate et al. (7) assuming a photobleaching time constant of *τ*_*pb*_ = 500 seconds. At this level of photobleaching the total fluorescent signal is attenuated by 73% by the end of the chase phase of the experiment. Panels A and B of Fig. 7 illustrate the state kinetics and fluorescent label kinetics predicted by the model for this fit to the data with *τ*_*pb*_ = 500 seconds. The predicted steady-state SRX fraction associated with this fit is 51%, significantly greater than the 34% level obtained assuming no photobleaching. Panel C shows the predicted SRX fraction (the steady-state value of 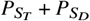) that is obtained for different values of *τ*_*pb*_. Controlling for photobleaching is thus an important factor to consider for quantifying the SRX properly. We do not have a way of estimating the actual effects of photobleaching on the data from the experiments analyzed here.

**Figure 7.**
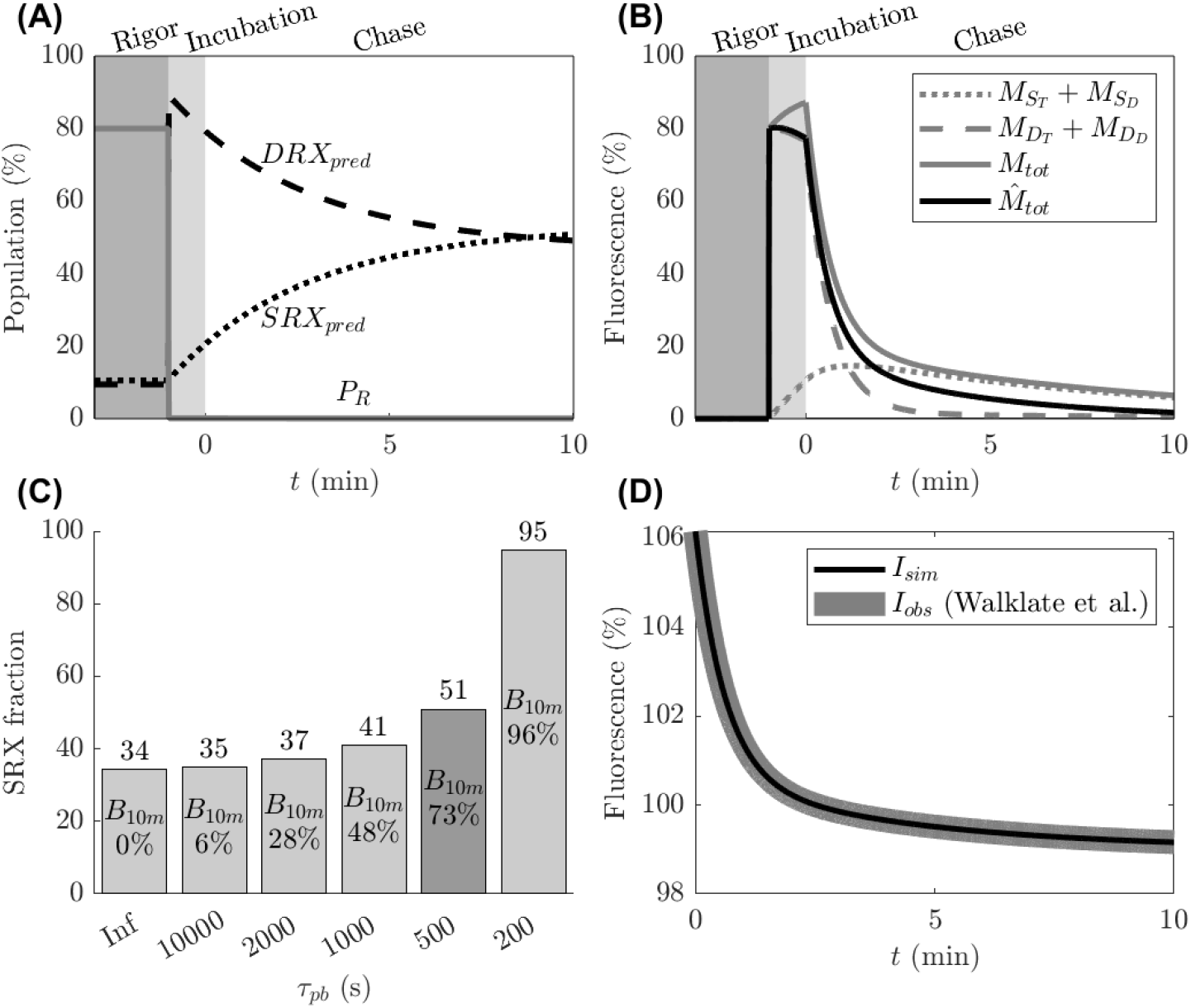
Effect of accounting for photobleaching on predicted SRX fraction, using *τ*_*pb*_ = 500s. (A) The simulated state probabilities show transition from the ratcheted (*R*) and ADP-bound (*P*_*D*_) states to the SRX (*P*_*S*_) and ATP-bound DRX 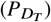 states (B) The predicted kinetics of fluorescent labeled states, with simulated photobleaching effect on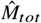. (C) Different values of *τ*_*pb*_ yield different predicted steady-state SRX fractions, with *B*_10m_ indicating the relative attenuation of fluorescence at 10 minutes. (D) Simulated fluorescent intensity *I*_*tot*_ during the chase phase is fit to reproduce the fluorescence decay in the ATP chase phase *I*_*obs*_ observed by (7) Fig. 1A using *τ*_*pb*_ = 500 seconds.

## DISCUSSION

We present a kinetic model of the myosin cross-bridge cycle and mant-ATP/ADP labeling to analyze the observed kinetics of mant fluorescence in mant-ATP label/chase assays and the biochemically defined SRX state of myosin. The model not only aligns with the available experimental data, but also provides a self-consistent explanation of three paradoxes summarized by Mohran et al. (6). Namely, the model: (1) predicts a steady-state balance between the states that is compatible with myosin cross-bridge models, (2) demonstrates discrepancy between experimentally estimated SRX fraction using double exponential fit and the simulated steady-state SRX fraction, and (3) offers a novel interpretation of the experiment of Walklate et al. (7) where the apparent SRX fraction varies little over experiments using incubation times from 0.2 to 30 seconds.

### Limitations of current mant-ATP assay interpretation

Contemporary conceptual models of mant-ATP decay assume an independent exchange of ATP in both SRX and DRX states (2, 6). Moreover, these models assume that both of these states undergo hydrolyzing exchange. That is, mant-ATP in both SRX and DRX can hydrolyze into mant-ADP + Pi, which may be replaced during the chase phase with unlabeled ATP. Our model is based on a fundamentally different kinetic scheme (Fig. 1), which is more consistent with existing models of the cross-bridge cycle.

Our analysis using this modeling framework reveals several potential important caveats for interpreting of mant-ATP label/chase experimental data:

1. Estimation of the SRX state occupancy based on double exponential analysis of the chase phase data, using either Equation (1) or (2), does not necessarily quantify the true fraction of the SRX state in steady state.
2. The SRX state is predicted to form relatively slowly (on the timescale of 5-10 minutes) during the incubation phase of a relaxed myosin. Since predicted time for the SRX and DRX states to reach steady-state is significantly longer than commonly used incubation times, transition into the SRX state continues even after the fluorescence intensity is saturated (Fig. 2). The use of relatively short incubation times leads to significant underestimation of the true SRX state based on the quantification of the slow decay component of the chase phase time course.
3. Photobleaching may substantially affect the observed decay and interpretation, resulting in underestimating the true SRX of the muscle (Fig. 7).

The uncertainties associated with these caveats may be addressed through the experimental design and the data analysis approach. Most fundamentally, it is important to track and account for experimental details that are often ignored, including the nature of the rigor incubation conditions, the incubation time, and the degree of photobleaching.

The fact that the double-exponential interpretation of mant-ATP data may not accurately quantify the true SRX fraction does not diminish its qualitative value. Under identical or comparable experimental conditions, it still provides insights into relative changes in the SRX population between samples. This study primarily emphasizes the limitations in absolute SRX quantification and the importance of interpreting such data with appropriate caution.

### Model configuration

Our model assumes a configuration in which the myosin may enter the SRX state only when ATP bound and that the SRX-bound ATP can undergo the same hydrolysis transition as DRX-bound ATP. Although transitions from *S*_*T*_ to *D*_*T*_ and from *D*_*D*_ to *S*_*D*_ cannot be ruled out, the double-exponential nature of the data provides only three effective degrees of freedom. As a result, more complex model configurations that introduce additional rate constants are not identifiable from the available data. Two alternative formulations, in which either the *S*_*D*_ or the *S*_*T*_ state is eliminated, are illustrated in Fig. 8. These model configurations were also evaluated and found to fit the data equally well. While detailed analyses of these models is out of the scope of this paper, the fits are accessible in the source code.

**Figure 8.**
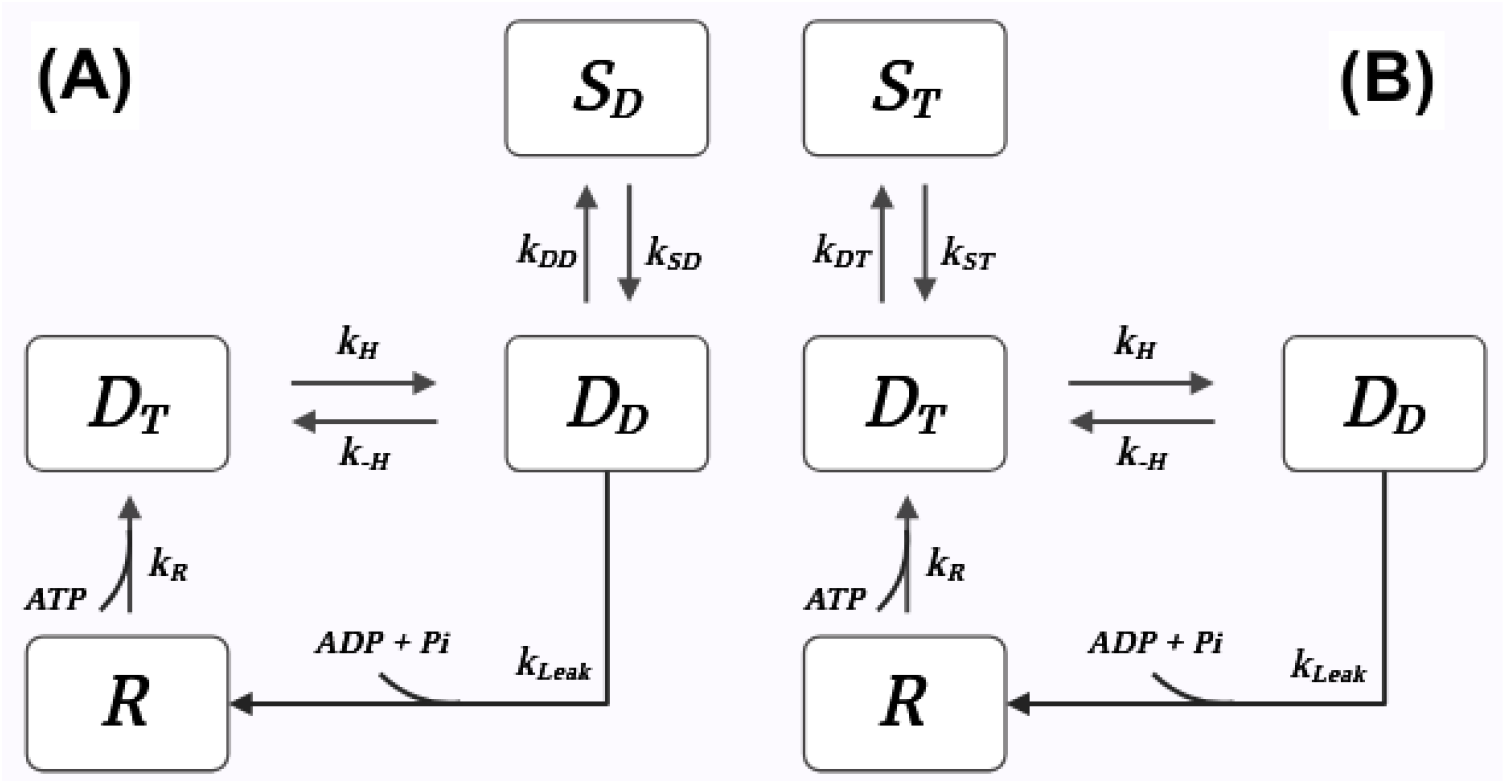
Alternative conceptual models. A) The myosin can form the SRX state only after ATP hydrolysis B) The SRX is formed and held while ATP bound only. Both of these conceptual models are able to fit the data equally well, with only minor differences in predicted SRX to the presented model.

### Model Limitations

The model has been tested on two published datasets. Although it fits these datasets very well, additional experiments under controlled conditions are necessary to validate the model’s predictions. The formulation of the model ignores potentially important effects, including diffusion (recently elaborated by Montesel et al (18)), SRX recruitment affected by the muscle tone ((13)), temperature effects, non-specific mant-binding, measurement artifacts (e.g. background fluorescence, microscope focus) and potentially many others (see e.g. (19)). While our model does not distinguish between blocked and free head myosins, there is some evidence of intermediate states of the heads of a myosin dimer entering the SRX state independently and the interaction of both heads and the tail stabilizing the SRX/IHM (20) but attempting to include these distinctions would only add significant complexity without clear experimental validation of the additional distinct steps. Also, the SRX might be unevenly distributed along the sarcomere length and stabilized by cMyBP-C((21–23)), a phenomenon incompatible with our probabilistic modeling approach. Interpretation of the model is valid only in context of the conditions simulated and caution should be exercised for comparing e.g. homogenized myosin protein (where e.g. atypic head conformations can exist) with whole tissue (prone to limitations caused by diffusion). Please note the SRX transition rates from the mant-ATP assay in this study (Table 2) are not directly comparable to values reported in the literature (e.g., (24, 25)), which are several orders of magnitude faster. This difference is mainly due to use of a different conceptual model, which includes the previously ignored *D*_*T*_ state. Second, additional considerations are required to apply the proposed model to active contraction. The muscle here is considered not activated and unloaded and thus the strong effects of force on SRX dynamics ((13, 26–28)) are not considered.

Moreover, dissociation and rate constants associated with mant-ATP/ADP binding and mant-APT/ADP associated states are assumed to be equal to kinetic constants associated with unlabeled ATP and ADP. However, Woodward et al.(29) report an approximately 10% decrease in mant-ATP hydrolysis rate (*k*_*H*_) and an approximately three fold faster mant-ATP binding (29) compared to the rates associated with unlabeled ATP hydrolysis and binding. Conflicting results exist regarding the mant-ADP affinity; Woodward et al. (29) report a 10-fold greater affinity of mant-ADP compared to unlabeled ADP, whereas (30) report no difference. Some of the effects are preliminary evaluated in the available source code, but given the uncertainty they are not accounted for in the presented model.

Although the model is able to essentially exactly match the time course decay data from the Walklate et al. and Hooijman et al. experiments, it does not capture the trend of decreasing apparent slow phase fraction with increasing incubation time observed by Walklate et al. (7). This discrepancy may reflect the presence of biochemical factors or mechanisms not accounted for in the model. However, the observed decrease in variability in the data with longer incubation aligns with model predictions of increased label accumulation during the incubation phase (Fig. 4B). A lower amount of bound label at shorter incubation times implies a lower signal-to-noise ratio, which could contribute to higher measurement uncertainty.

## CONCLUSION

Our analysis suggests that apparent paradoxical interpretations of the mant-ATP label/chase experiment to assess myosin states may be explained using a kinetic model, while accounting for potential protocol differences.

Application of this kinetic framework may improve the quantification and interpretation of the label/chase experiment. For example, we estimated the actual SRX fraction to be 34% in a representative stopped-flow experiment (Walklate et al., 2022), compared to the originally reported 23%, based on the slow-phase fraction double-exponential fit. The presented model may be used for enhanced quantification of the SRX state occupancy and investigation into the transition rates.

## AUTHOR CONTRIBUTIONS

FJ, SJH, ASVR and DAB conceptualized the research, ASVR supervised the project and validated the outcomes. FJ and SJH developed the model and performed simulations, FJ, SJH, ASVR and DAB analyzed and interpreted the results. FJ and DAB drafted the manuscript, SJH and ASVR edited the manuscript.

## ACKNOWLEDGMENTS

We appreciate a thoughtful discussion with Kenneth Campbell.

This work was supported by National Heart, Lung and Blood Institute (NHLBI) Grants R01 HL154624 (D.A.B.), R01 HL173346 (D.A.B.) and R00 HL153679 (A.S.V.R.) and National Institute of General Medical Sciences (NIGMS) RM1 GM131981 (A.S.V.R).

This manuscript is the result of funding in whole or in part by the National Institutes of Health (NIH). It is subject to the NIH Public Access Policy. Through acceptance of this federal funding, NIH has been given a right to make this manuscript publicly available in PubMed Central upon the Official Date of Publication, as defined by NIH.

## SUPPLEMENTARY MATERIAL

Codes and data supporting this paper are published at https://github.com/beards-lab/mantATP-SRX.

